# Organization of cortico-hippocampal networks in the human brain

**DOI:** 10.1101/2020.06.09.142166

**Authors:** Alexander J. Barnett, Walter Reilly, Halle R. Dimsdale-Zucker, Eda Mizrak, Zachariah Reagh, Charan Ranganath

**Author notes:** Corresponding Author Alexander J. Barnett, 1544 Newton Ct, Davis, CA 95618, United States Center for Neuroscience, Room 209. Author Contributions: AJB, WR, HDZ, and CR designed research; AJB, HDZ, EM, and ZR performed research; EM contributed new reagents or analytic tools; AJB and WR analyzed data. AJB and CR wrote the paper.

## Abstract

Episodic memory depends on interactions between the hippocampus and the interconnected regions comprising default mode network (DMN). Here, using data-driven analyses of resting-state fMRI data, we identified the networks that interact with the hippocampus—the DMN and a “Medial Temporal Network” (MTN) that included regions in the medial temporal lobe and retrosplenial cortex. We observed that the MTN plays a critical role in connecting the visual network to the DMN and hippocampus. The DMN could be further divided into three subnetworks: a “Posterior-Medial” Subnetwork comprised of posterior cingulate, and lateral parietal cortices, an “Anterior-Temporal” Subnetwork comprised of regions in the temporopolar, and dorsomedial prefrontal cortex, and a “Medial-Prefrontal” Subnetwork comprised of regions primarily in the medial prefrontal cortex. These networks vary in their functional connectivity along the hippocampal long-axis and represent different kinds of information during memory-guided decision-making. Finally, a Neurosynth meta-analysis of fMRI studies suggests new hypotheses regarding the functions of the MTN and DMN subnetworks, providing a framework to guide future research on the neural architecture of episodic memory.

---

Episodic memory allows us to relive past events that happened at a particular place and time (1). Most research investigating the neurobiology of episodic memory retrieval has focused on the hippocampus and medial temporal lobe (MTL) cortex (perirhinal (PRC), parahippocampal (PHC), and entorhinal cortex) highlighting the interactions among these regions (2–4). It is generally assumed, however, that the hippocampus supports memory by integrating information represented across distributed areas in the neocortex (5). Consistent with this idea, fMRI studies of memory retrieval have shown activity and connectivity in a cortico-hippocampal network composed of medial parietal, lateral parietal, lateral temporal, and medial prefrontal neocortical regions along with the MTL (6–10), and lesion studies have shown that damage within this network can cause amnesia (11).

This distributed set of cortical regions that are recruited during episodic retrieval overlaps extensively with regions that comprise the default-mode network (DMN). The DMN is a large-scale network that is typically identified in studies that use intrinsic functional connectivity analysis of functional magnetic resonance imaging (fMRI) data (12), and some studies suggest that the DMN can be differentiated into different subnetworks (13–15). One approach to partition the DMN has been to view DMN subnetworks as an extension of the connectivity differences within the MTL (16–18). Guided by neuroanatomical studies of MTL connectivity in rodents and nonhuman primates (19–22), studies of intrinsic functional connectivity in fMRI data (23–26) have differentiated between cortical regions that preferentially affiliate with the PRC and anterior hippocampus and cortical regions that preferentially affiliate with the PHC and posterior hippocampus (27, 28). For example, Libby et al. (24) demonstrated that the PHC has relatively higher functional connectivity with a network of Posterior Medial (PM) cortical regions, including the posterior cingulate and retrosplenial cortex and posterior hippocampus, whereas the

PRC has relatively higher connectivity with a network of Anterior Temporal (AT) cortical regions, including the orbitofrontal and temporopolar cortex and anterior hippocampus. Drawing on these findings, Ranganath and Ritchey (29) reviewed evidence converging on the idea that PM and AT networks support different aspects of memory-guided behavior. This “PM/AT framework” has provided a valuable framework for interpreting memory phenomena in imaging (9, 30–32), stimulation (33, 34), and disease (35–37). However, recent work has come to question the homogeneity of the PM network (38), with several studies highlighting a dissociation between PHC, retrosplenial and precuneal cortex from posterior cingulate cortex (PCC) (39, 40) under conditions that seem to tax perceptual relative to abstract event processing, or scene relative to face processing.

A second approach to segmenting the DMN has used data-driven methods and these studies have also found partially conflicting network delineations (15, 41, 42). These studies have generally partitioned the DMN into three subnetworks: an MTL subnetwork, a midline subnetwork of posterior cingulate and medial prefrontal cortex, and a third network composed of dorsal medial prefrontal, lateral temporal, and ventrolateral prefrontal cortex, but have focussed largely on cortical regions. Critically, it is unclear how these data-driven, cortical networks pertain to memory functioning and previous frameworks that have strongly characterized hippocampal connections.

Here, we seek to unify these two approaches. The MTL-centric approach has rarely considered the interplay of regions outside of the MTL, and a recent review has suggested that considerable heterogeneity may exist within the previously described PM network (38), prompting a need for re-examination of these network properties. Conversely, the data-driven approach has rarely described the hippocampal interactions with these networks, focusing mainly on cortico-cortical interactions limiting the applicability to understanding episodic memory. The goal of this paper was to comprehensively characterize cortico-hippocampal networks that contribute to episodic memory. Using a whole-brain, data-driven approach to examine functional connectivity in resting-state fMRI, we sought to 1. identify and partition the DMN into subnetworks and examine whether these subnetworks converge with the PM/AT framework, 2. examine the connectivity of the hippocampus to the identified networks, and 3. determine whether these cortico-hippocampal networks play different roles in episodic memory.

## Results

### Data-Driven Partition of Neocortical Networks

Our first objective was to use a data-driven approach to identify large-scale resting-state networks. Using 25 minutes of resting-state fMRI data acquired from 40 participants, we partitioned the brain into canonical resting networks using state-of-the-art techniques (43). We extracted the average confound-corrected timeseries from each region in a recently published atlas of the human neocortex (44) and computed Fisher z-transformed Pearson’s correlations between the timeseries of each cortical region and every other cortical region. We then created a group-averaged functional connectivity matrix to be used for community detection. The Louvain community detection algorithm (45) was run for 1000 iterations, tuning the resolution (how large or small communities might be) across a range of resolution parameters to identify a partition solution that showed both high network modularity (i.e. higher within community connectivity than would be expected by chance) and high network grouping consistency (quantified using the z-Rand index (43, 46)). The solution was also subject to a qualitative criterion that the partition should separate the primary sensory networks (43). The resolution parameter of gamma = 2.005 satisfied our criteria and produced a network partition (Figure 1) that corresponds closely with previously reported partitions in other recent studies (43, 47, 48). This partition identified the DMN, but, interestingly, some areas (such as the PHC, retrosplenial cortex, and PRC) that are often identified as part of the DMN were grouped within a separate network that included MTL and parietal regions. This set of regions overlaps with what has been previously described as the MTL network (13), or the contextual association network (49), and coactivates particularly during recall of places (50, 51). Here we will describe this network as the medial temporal network (MTN).

**Figure 1.**
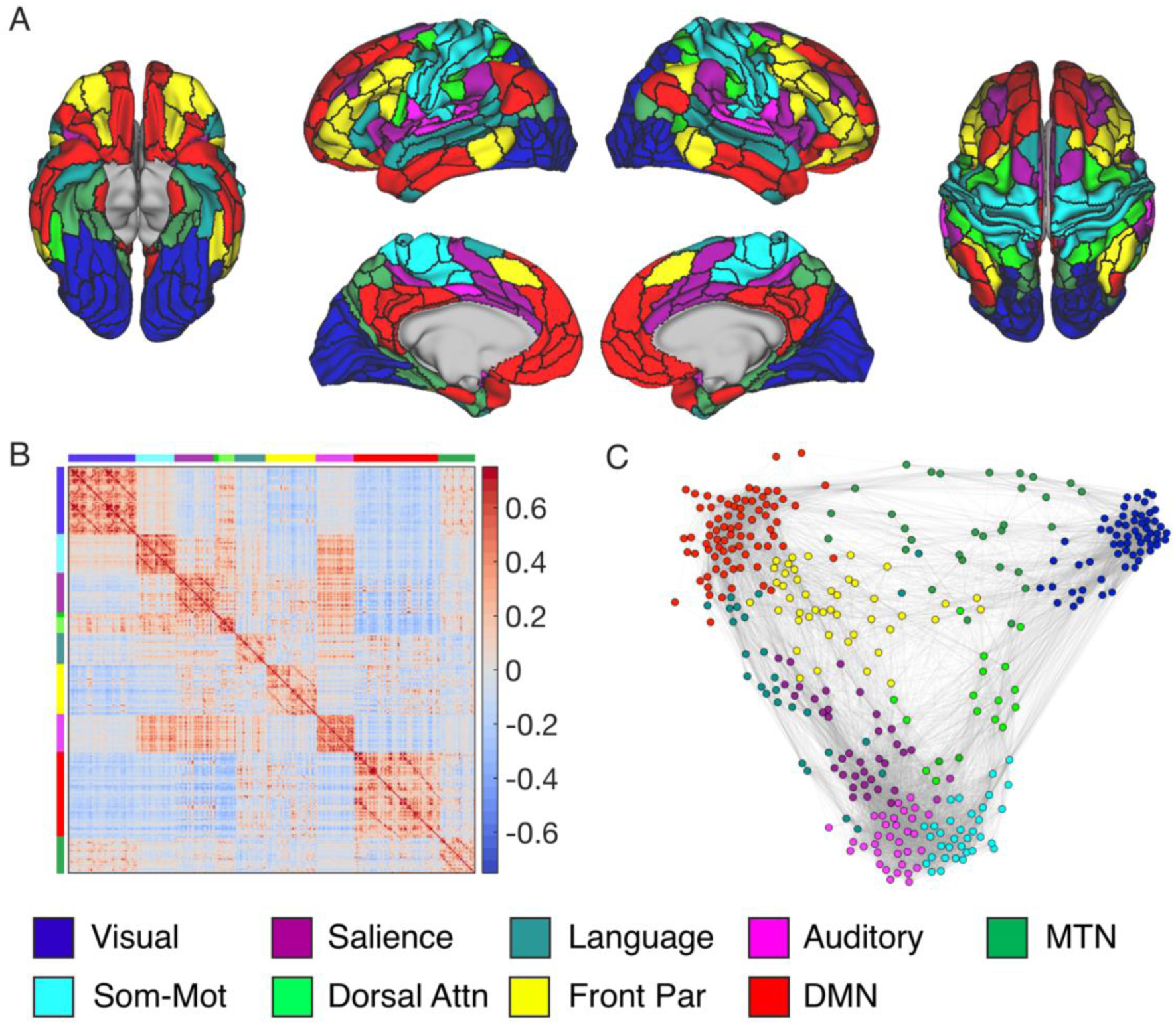
Louvain community detection identifies large-scale resting-state networks including the default-mode and medial temporal networks that are thought to interface with the hippocampus. A. Inflated cortical surface, colored according to community membership. B. Connectivity matrix reordered by community to demonstrate the community structure of the group-averaged brain. Colors along the axis demonstrate which rows/columns belong to a given community. Color bar represents Fisher Z-transformed correlation values. C. Force-directed graph of the group-averaged networks color coded by community membership of the selected partition created using the ForceAtlas2 algorithm (52). Attn, attention; DMN, Default mode network; Front Par, Frontoparietal; MTN, Medial temporal network; Som-Mot, Somatomotor.

We next interrogated the functional connectivity between the networks defined above and the hippocampus. We segmented the hippocampus in each participant using FreeSurfer v6.0 (http://surfer.nmr.mgh.harvard.edu/, (53)), and then manually segmented it further into anterior and posterior regions, as these subdivisions of the hippocampus are known to have somewhat different functional properties (16, 20). The posterior hippocampus was defined as all the hippocampus posterior to the last slice of the uncal notch (16). We calculated the connectivity of the hippocampus to every cortical region in the atlas and averaged together functional connectivity weights of cortical regions within the same network based on network affiliations. This was done for each participant. One-sample t-tests revealed that two networks were functionally connected to every hippocampal ROI in our sample—the DMN and the MTN (DMN range: *t*(38) = 10.7–12.9, *all P* < .001; MTN range: *t*(38) = 4.2–9.3, *all P* < .01). The language network and somatomotor network both showed mixed connectivity with the hippocampus, showing significant anterior, but not posterior hippocampal connectivity (see Supplemental Table 1). The robust connectivity of the MTN and DMN to the hippocampus corroborates previous investigations of hippocampal connectivity (54).

### Network analysis support an interfacing role of the MTN

A recent review discussing the heterogeneity within the PM network suggested that parahippocampal, and lateral parietal cortex interface with lower level sensory cortex in service of higher-order feature encoding prior to binding in the hippocampus (29, 38). They also note that these regions also interface with other DMN regions such as the mPFC and PCC (regions found in the DMN, here) that track long-timescale event structure and provide conceptual knowledge pertaining to event schemas (38, 55). Thus, the MTN may serve as a bridge between external, low level perception in the visual network and the hippocampus and DMN. By plotting the structure of the network interactions in Figure 2A (left) we do indeed see that the MTN shows functional connectivity with both the DMN, hippocampus, and the visual network. Treating the brain as an abstract graph (56), we sought to test the hypothesis that the MTN mediates network communication between the visual network and the DMN and hippocampus. To do so, we used graph theory to calculate the shortest path length between the visual network and the DMN and hippocampus. Path length refers to the shortest number of links that needs to be traversed to link one node in a network to a target node. If two regions are directly connected by a strong link, they will have a short path length. If two regions are not directly connected by a positive link but can be connected by multiple intermediate nodes that are positively connected, then they would have a longer path length. To examine the influence of the MTN on internetwork communication between the DMN and visual network, we compared the path length between the visual network and DMN when the MTN was removed relative to the path length when we removed all other networks. If the MTN mediates the connectivity between the visual network and DMN and hippocampus, then removal of the MTN should disproportionately increase the average path length between visual network and DMN over and above any path length increases caused by removal of all other networks. This was done at the individual-subject level to determine the significance of the effect and is similar to methods used by (57).

**Figure 2.**
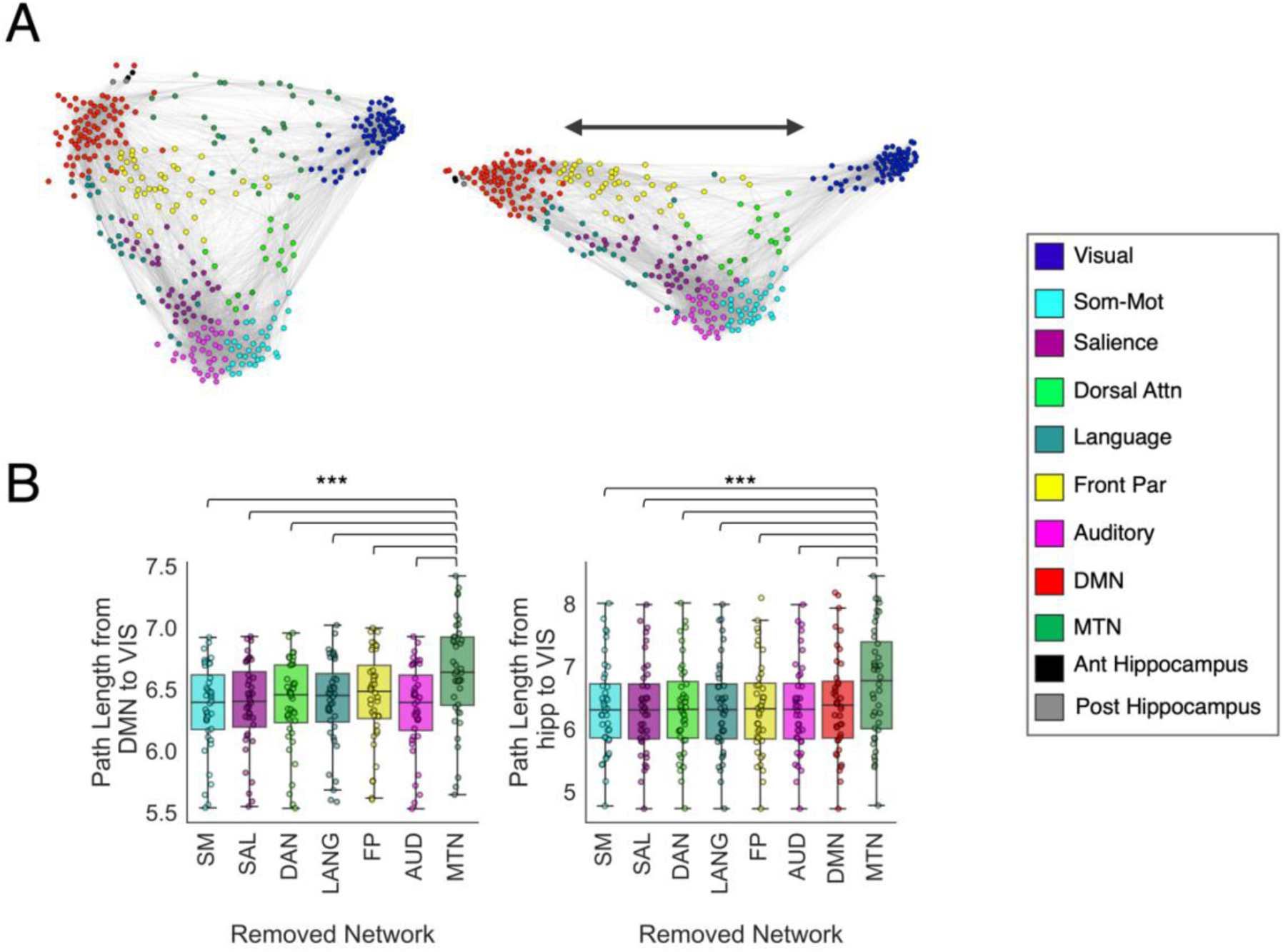
Removal of the MTN disproportionately decreases the network connectivity of the DMN and visual network. (A) Force-directed graph of the group-averaged networks color coded by community membership of the selected partition created using the ForceAtlas2 algorithm (52) with the MTN present (left) and with the MTN excluded (right). (B) Boxplots displaying the path length between the DMN and visual network or the path length between the hippocampi and visual network following removal of each network. *** p < .001.

We found that across subjects, removal of the MTN led to a disproportionately large increase in DMN–visual network path length compared to removal of any other network (all *t*(39) > 6.7, p < .001; Figure 2B), indicating that the MTN plays a critical role in cross-network communication between the visual network and DMN. Importantly, this effect cannot be explained simply by the number of nodes removed from the whole network, as the MTN has fewer nodes (30 nodes) than the somatomotor (34 nodes), salience (35 nodes), frontoparietal (44 nodes) and auditory (33 nodes) networks. To further confirm this, we randomly removed 30 nodes from each subject’s network (excluding DMN, and visual nodes) and recalculated the path length values. This was performed 1000 times and we compared the DMN-visual network path length of the true removal of the MTN nodes compared to the distribution of DMN-visual network path length following random node removals. We observed that the MTN removal still resulted in greater average path length between the DMN and visual network relative to removal of random nodes, *t*(39) = 9.4, *p* < .001. Another possible explanation is that due to the high level of connectivity between the DMN and MTN, removal of the MTN will non-specifically lead to a disproportionate increase in path length of the DMN to all other networks. However, the path length effects caused by removal of the MTN were specific to the DMN – visual network path lengths (S1 Figure).

As mentioned above, it is well-known that MTL regions such as the perirhinal and parahippocampal cortex provide the hippocampus with higher-order representations of objects and scenes (4, 58). Therefore, we repeated the previous analysis, this time examining the effects of MTN removal on the pathlength between the hippocampus and visual network. We observed that removal of the MTN lead to greater path length between the hippocampus and visual network compared to removal of any other networks (all *t*(39) > 6, p < .001; Figure 2B). Again, MTN removal led to disproportionately higher path length than removal of random nodes (*t*(39) = 3.15, *p* = .003). This effect was specifically between the hippocampus and visual network path length, rather than leading to disproportionately higher path length between the hippocampus and any other network, S2 Figure.

### Partitioning the DMN

We next sought to identify whether the DMN could be broken up into subnetworks using similar methods that we used in our whole-brain network partition. We selected the DMN regions identified in our whole-brain partition and performed Louvain community detection on the functional connections between the regions in the network. This was done for 1000 iterations across a range of resolution parameters, and we again selected the partition yielding the highest modularity and stability. The solution with the highest modularity-weighted stability produced a partition of the DMN with three communities which we refer to as the PM subnetwork, the AT subnetwork and the medial prefrontal (MP) subnetwork. The PM subnetwork and AT subnetwork were named as such based on their resemblance to previous spatial maps of these networks (24). The PM subnetwork included posterior cingulate cortex, posterior angular gyrus, and dorsal prefrontal cortex. The AT subnetwork included temporopolar cortex, lateral orbitofrontal cortex, temporoparietal junction and dorsomedial prefrontal cortex. The MP subnetwork included medial prefrontal regions and the entorhinal cortex (Figure 3).

**Figure 3.**
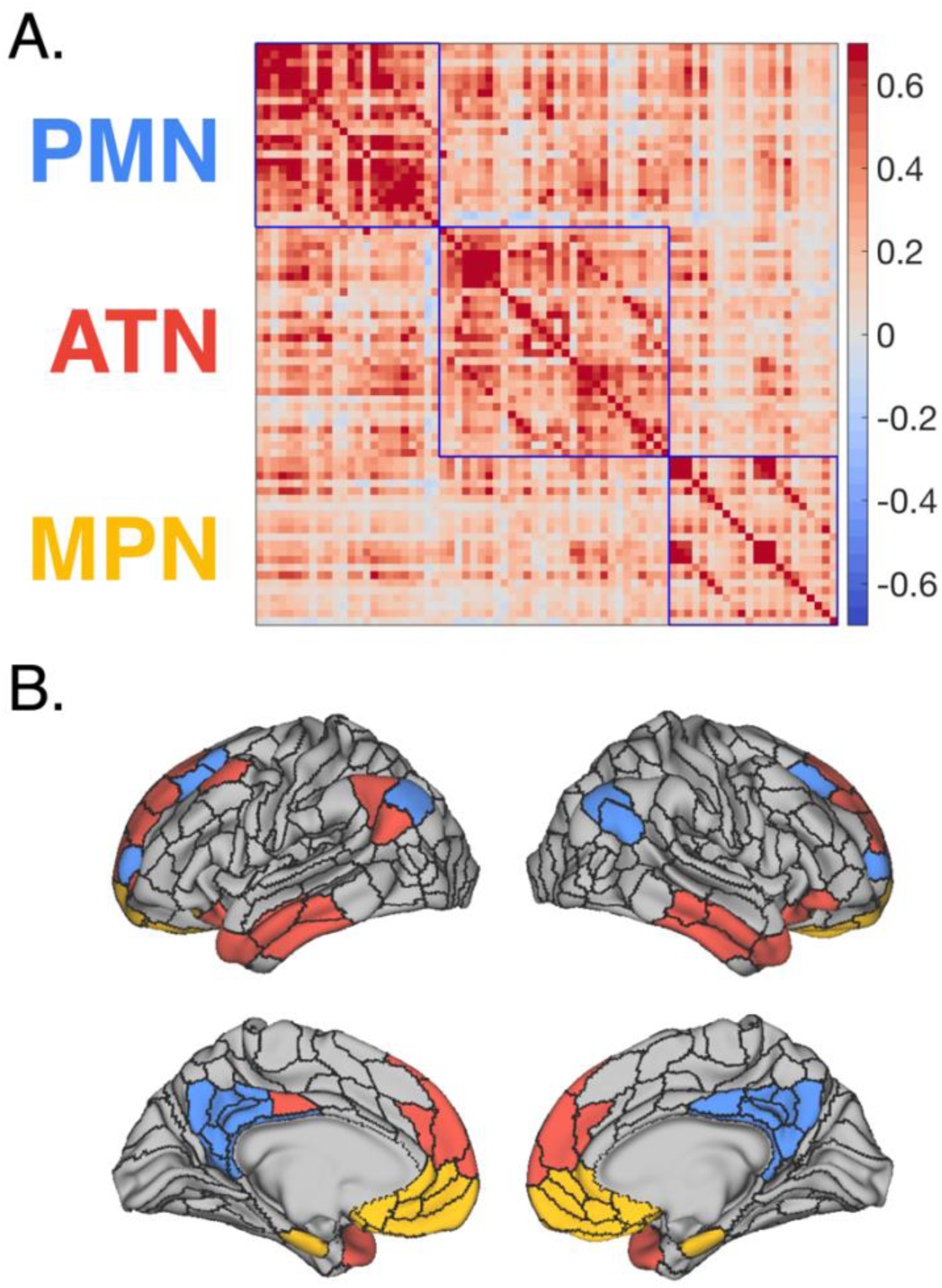
The default-mode network can be partitioned into three interconnect subnetworks. A. Functional connectivity matrix organized by subnetwork. Color bar represents Fisher Z-transformed correlation values. B. Inflated cortical surface, colored according to community membership of subnetworks. AT, anterior temporal; MP, medial prefrontal; PM, posterior medial.

To visualize the topology of connections between these DMN sub-networks and the other large-scale networks defined earlier, we constructed a force-directed graph in abstract graph space (Figure 4). The ForceAtlas2 algorithm was used to construct the layout by modeling the graph as a physical system in which nodes repel each other and the functional connectivity between the nodes act as springs (52). Regions within the PM subnetwork tightly clustered and several nodes showed prominent cross-network connectivity with the MTN and frontoparietal network. The AT subnetwork showed cross-network interactions with the language and frontoparietal network and the MP subnetwork was less clustered with fewer out of DMN connections.

**Figure 4.**
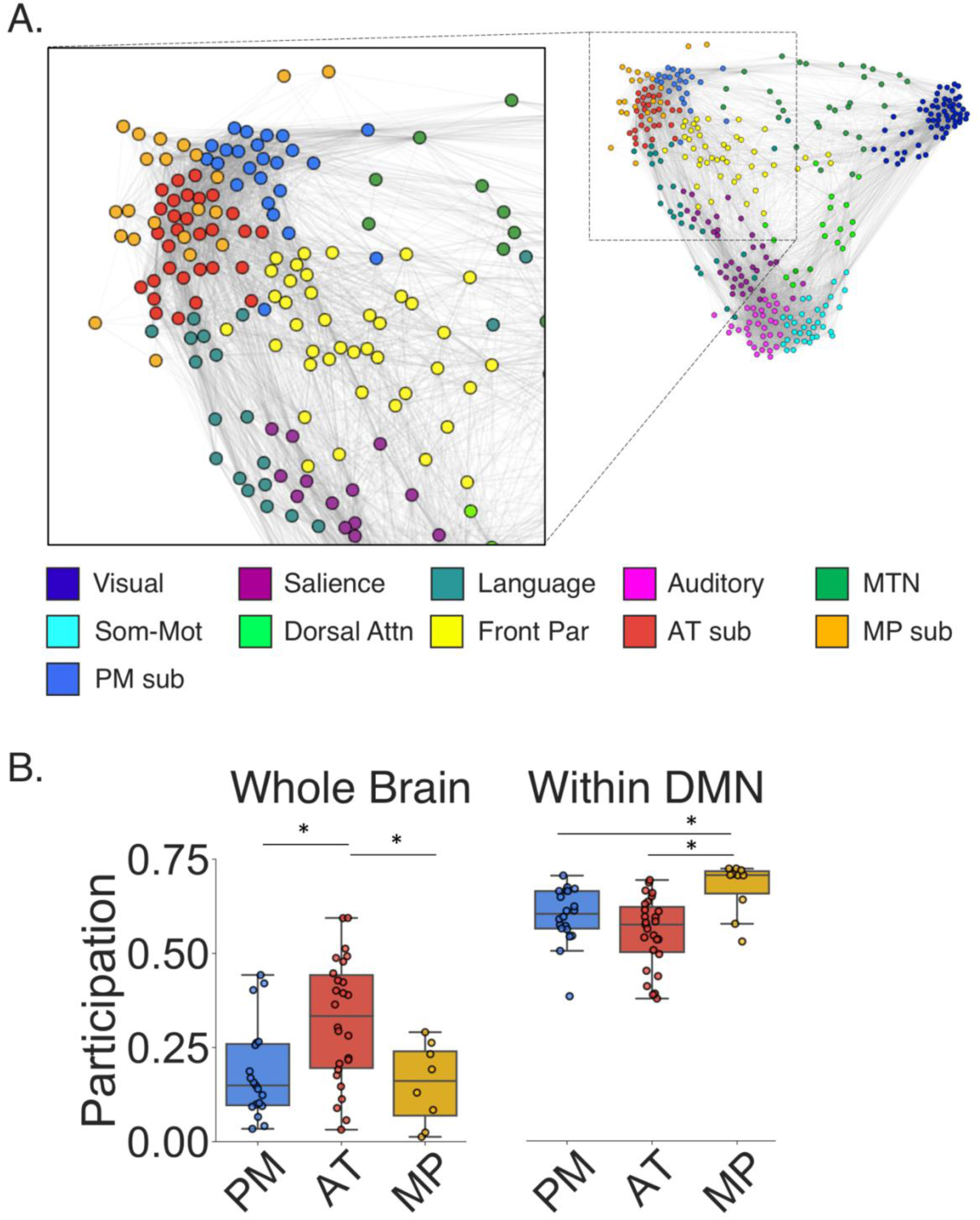
The subnetworks of the DMN interact with language, frontoparietal and medial temporal networks. A. Force-directed graph of the group-averaged networks with the DMN relabelled according to subnetwork membership. B. Boxplots showing participation coefficient for DMN nodes to the rest of the brain (Whole Brain) and participation coefficient for DMN nodes to subnetworks of the DMN (Within DMN). AT, anterior temporal; Attn, attention; Front Par, Frontoparietal; MP, medial prefrontal; MTN, medial temporal network; PM, posterior medial; Som-Mot, Somatomotor. *indicates significant difference at p < .05.

To quantify the diversity of connections in the DMN subnetworks to the rest of the brain, we calculated participation coefficients—a graph theory metric that describes the extent to which a node’s functional connections are spread across different communities. Nodes with a high participation coefficient are thought to be critical for communication between communities, and disruption of such nodes can lead to altered network function and cognitive impairment (59–62). Of the subnetworks, the AT subnetwork had the greatest diversity of connections to nodes outside of the DMN (AT > PM: *t*(43) = 3*, P* =.004; AT > MP: *t*(32) = 2.7*, P* =.01). As shown in Figure 4B, the PM subnetwork had several nodes with relatively high participation. These nodes connect to MTN, and the frontoparietal network, but the majority of the PM subnetwork showed less out of network connectivity. The MP subnetwork, on the other hand, had relatively low participation outside of the DMN.

We next addressed the extent to which nodes in the DMN showed diverse connectivity with nodes in the other DMN subnetworks. Interestingly, regions in the MP subnetwork exhibited the highest participation coefficients across the DMN subnetworks (MP > PM: *t*(28) = 2.7*, P* = .01; MP > AT: *t*(35) = 3.7*, P* < .0007). These findings substantiate what can be seen in Figure 4A—that nodes in the MP subnetwork are positioned to integrate information across DMN subnetworks, whereas nodes in the AT subnetwork and PM subnetwork may interface with networks outside of the DMN.

### Cortico-hippocampal network connectivity

Having identified cortical networks and subnetworks that connect to the hippocampus (cortico-hippocampal networks), we then examined whether these networks showed a preference in connectivity along the hippocampal long-axis. Previous research has shown differential functional connectivity along the long-axis axis of the hippocampus (24, 28, 63, 64), with the anterior hippocampus demonstrating greater connectivity to orbitofrontal cortex, and temporal pole and the posterior hippocampus demonstrating greater connectivity to retrosplenial cortex and precuneus. We therefore hypothesized that the anterior hippocampus should have preferential connectivity to the AT subnetwork, whereas the posterior hippocampus would have preferred connectivity with the PM subnetwork. These predictions were partially confirmed by our findings. The anterior hippocampus showed stronger connectivity to the AT and MP subnetworks *(t(*37) = 8.2*, P* < .00001; *t*(37) = 6.5*, P* < .00001) and the posterior hippocampus showed stronger connectivity to the MTN than the anterior hippocampus *(t(*37) = 2.6*, P* = .01), but there was no significant difference in connectivity between the hippocampal regions and PM subnetwork (*t*(37) = .41, *P* = .68) (Figure 5A). To determine whether these differences were evenly spread across the networks or driven by particular regions within the networks, we contrasted anterior and posterior hippocampal connectivity at a region-to-region level. As expected from the network-level analysis, we observed significantly greater anterior hippocampal connectivity to medial prefrontal, anterior temporal, and also anterior medial temporal cortex, whereas the posterior hippocampus had significantly greater connectivity to the parietal occipital sulcus, precuneus, and dorsal posterior cingulate cortex. Thus, the anterior-posterior differences are fairly consistent in the AT and MP subnetworks, but within the MTN the overall anterior-posterior differences are driven by relatively higher posterior hippocampal connectivity in the medial parietal cortex (Figure 5B).

**Figure 5.**
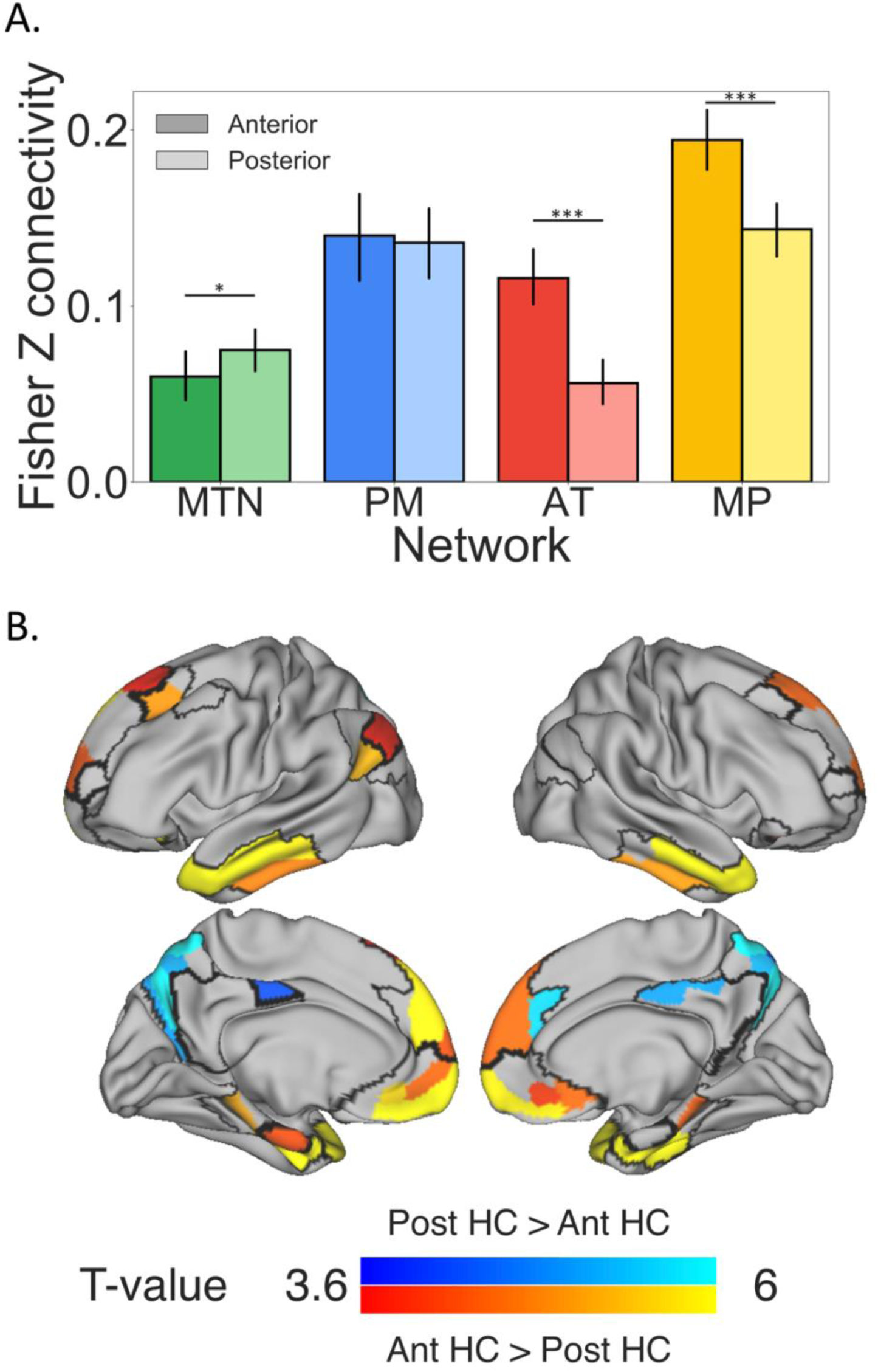
Regions within the MTN and DMN subnetworks exhibit differential functional connectivity with the anterior and posterior hippocampus. A: Bar graphs of functional connectivity from each of the cortical networks to the anterior and posterior hippocampus. Error bars show 95% confidence interval. B: T-contrast of anterior hippocampal connectivity versus posterior hippocampal connectivity, with warm colors showing regions that had stronger anterior hippocampal connectivity and cool colors showing regions that had stronger posterior hippocampal connectivity. Cortico-hippocampal network boundaries are outlined on the surface of the brain. These results indicate that the temporopolar and medial prefrontal cortex show preferential connectivity with the anterior hippocampus, whereas posterior medial parietal cortex shows preferential connectivity with the posterior hippocampus. Ant, anterior; AT, anterior temporal; HC, hippocampus; MP, medial prefrontal; MTN, medial temporal network; PM, posterior medial; Post, posterior. *\* P* < .05, \**\*\* P* < .001.

Based on neuropsychological evidence demonstrating distinct functional roles of the left and right hippocampus (65–67), we next investigated whether the left and right hippocampus differed in their connectivity to the cortical networks. We observed significant hemispheric laterality effects with the right hippocampus having greater connectivity to the MTN than the left (*t*(37) = 3.8, *P* = .0001) and the left hippocampus having greater connectivity to AT subnetwork (*t*(37) = 2.6, *P* = .01). There was no significant laterality effect in the PM (*t*(37) = .29, *P* = .77) and MP subnetworks (*t*(37) = .98, *P* = .33). This connectivity difference might help to explain differentiated function of the left and right hippocampus.

### Regions within the same community represent similar kinds of information during a memory task

We next sought to determine whether network membership was related to functional relevance of these subnetworks using an independent dataset in which participants performed a memory retrieval task (described in Mizrak et al. (68) and in *Methods* section) by examining the representational profile similarity between regions within the same network or in different networks (17, 69). Multivoxel pattern similarity was calculated between trials which created a trial-by-trial representational similarity matrix for every ROI in our cortico-hippocampal networks. Here, we assume that trial-by-trial fluctuations in pattern similarity for a given ROI are driven by trial-by-trial fluctuations in features for which the ROI is sensitive. These fluctuations across trials represent the ROI’s *representational profile*. We sought to determine whether regions within the same network had greater similarity in their representational profiles—and thus represent similar features—compared to regions outside their network.

Here, we calculated the similarity of representational profiles between ROIs by correlating each ROI’s representational similarity matrix to every other ROI. We then averaged the representational profile similarities for pairs of ROIs within the same network to create a mean within-network similarity value, for each subject (we removed the similarity of each ROI with itself to avoid inflating the within network similarity). To create a mean between-network similarity value, for each subject, we averaged together the representational profile similarity values for pairings of ROIs that were in different networks. We contrasted the mean within-network similarity with the mean between-network similarity using a repeated-measures ANOVA and observed that regions within the same network had higher profile similarity compared to regions in different networks, *t*(21) = 18.5*, P* < .001 (Figure 6). This effect persists even when accounting for the similarity in trial-by-trial mean BOLD activity (*t*(21) = 12.1, p < .001), suggesting it is not simply a result of univariate activity similarity.

**Figure 6.**
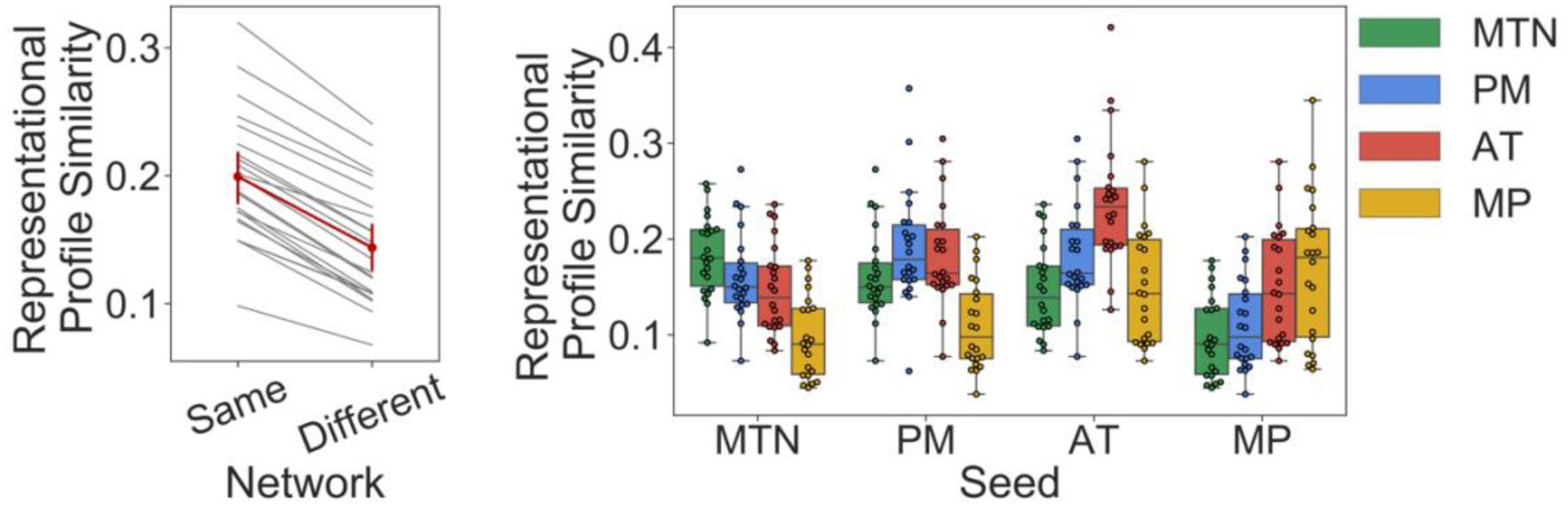
Regions show higher representational profile similarity with community members compared to regions in other communities. Left: Individual subject representational profile similarity of regions within the same network or between network in grey lines with the group average in red, with 95% confidence intervals represented by error bars. Right: Boxplot of average region-to-region representational profile similarity between each cortical-hippocampal network (labelled along the x-axis) to itself and every other cortico-hippocampal network (each represented by a colored bar) across the sample. Individual participants represented as dots. AT, anterior temporal; MP, medial prefrontal; MTN, medial temporal network; PM, posterior medial.

Another possible explanation of these findings is that regions in the same network tend to be closer in spatial proximity and there is some spatial non-independence of adjacent ROIs of these representational effects. To account for this spatial proximity, we created an ROI-by-ROI distance matrix by calculating the Euclidean distance between the center of gravity of all pairs of ROIs. We then regressed out the influence of spatial proximity from the representational profile matrix using this distance matrix and repeated the analysis. After statistically removing the influence of spatial proximity, we still observed that ROIs within the same network had greater representational profile similarity compared to pairs of ROIs in different networks, *t*(21) = 9.3, p < .001.

### Neurosynth Decoding

Having found evidence that regions within the same cortico-hippocampal networks carry similar information, we sought to identify the cognitive terms typically associated with activity in these networks. To better understand the role these networks may play in cognition, we used Neurosynth (70) to determine what cognitive terms are typically associated with activation in these subnetworks. The meta-analytic activation map of every term in the Neurosynth database was correlated with the binarized volumetric masks of each cortico-hippocampal network. By examining the top cognitive terms associated with each network we observed that the term autobiographical was the sole common term in the top 20 across all networks. Activation in MTN and PMN is disproportionately associated with studies involving episodic memory, but this was not the case for the ATN and MPN in which the term episodic memory did not appear in the top 20 search terms. The MTN also showed strong overlap with meta-analytic maps for preference for navigation (*r* = .15), scenes (*r* = .12), and the PM subnetwork showed strong overlap with self-referential *(r* = .13), and theory of mind terms *(r* = .08). The top terms associated with AT subnetwork related to theory of mind *(r* = .22), intentions *(r* = .17), mental states *(r* = .22), and social cognition *(r* = .19), while the top terms associated with MP subnetwork were value *(r* = .17), fear *(r* = .16), emotion *(r* = .16), and terms related to emotional valence *(r* = .14). Weightings for each network are available in Supplementary Table 2.

## Discussion

We have taken a data-driven approach using resting-state fMRI to identify and characterize cortico-hippocampal network connectivity, finding a set of subnetworks that interact with the hippocampus. Our analyses showed that the MTN, a cluster of regions in the medial temporal and dorsomedial parietal lobes, could be differentiated from the DMN, which included medial and lateral parietal, temporal, and frontal regions. This MTN mediates connectivity between the visual network and the DMN and between the visual network and hippocampus. We also found that the DMN could be meaningfully subdivided into three subnetworks: the PM subnetwork, encompassing the posterior cingulate, lateral parietal, and dorsal lateral prefrontal cortex, the AT subnetwork, encompassing the temporopolar, lateral orbitofrontal, and dorsal medial prefrontal cortex, and the MP subnetwork, encompassing the ventral medial prefrontal, and entorhinal cortex. These cortico-hippocampal subnetworks vary in connectivity strength along the long-axis of the hippocampus, and meaningfully segregate based on feature representation as measured by multivoxel activity patterns in an independent dataset.

Prior work on the neurobiology of memory has drawn on the idea that the hippocampal formation primarily interacts with the PHC and PRC, such that these areas collectively comprise a “MTL memory system” (2) that is functionally distinct from surrounding cortical areas. More recently, we and others proposed that PHC and PRC are embedded in larger scale cortico-hippocampal networks as evidenced by divergent anatomical pathways found in rodents and non-human primates (19–22) and by differing functional connectivity of the PHC and PRC in humans (24, 25, 71). In this “PM/AT” framework, the PHC is a core region in the PM network, which also includes retrosplenial, posterior cingulate and lateral parietal cortex, whereas the PRC is a core region in the AT network, which also includes temporopolar, and orbitofrontal cortex (29). As shown in Figure 8, the present results were not fully consistent with either the original PMAT framework. We found that the PRC, PHC, retrosplenial cortex, inferior lateral parietal, and dorsal medial parietal formed the MTN, and this network could be distinguished from the DMN. This network corresponds closely to the “parieto-medial temporal pathway” described by Kravitz et al. (72). This network division converges with previous data-driven community detection studies (15, 42, 47, 73) showing that the MTN can be differentiated from the DMN and support recent suggestions that heterogeneity exists within the PM system (38, 50). Critically, both the MTN and DMN showed strong functional connectivity with the hippocampus.

**Figure 7.**
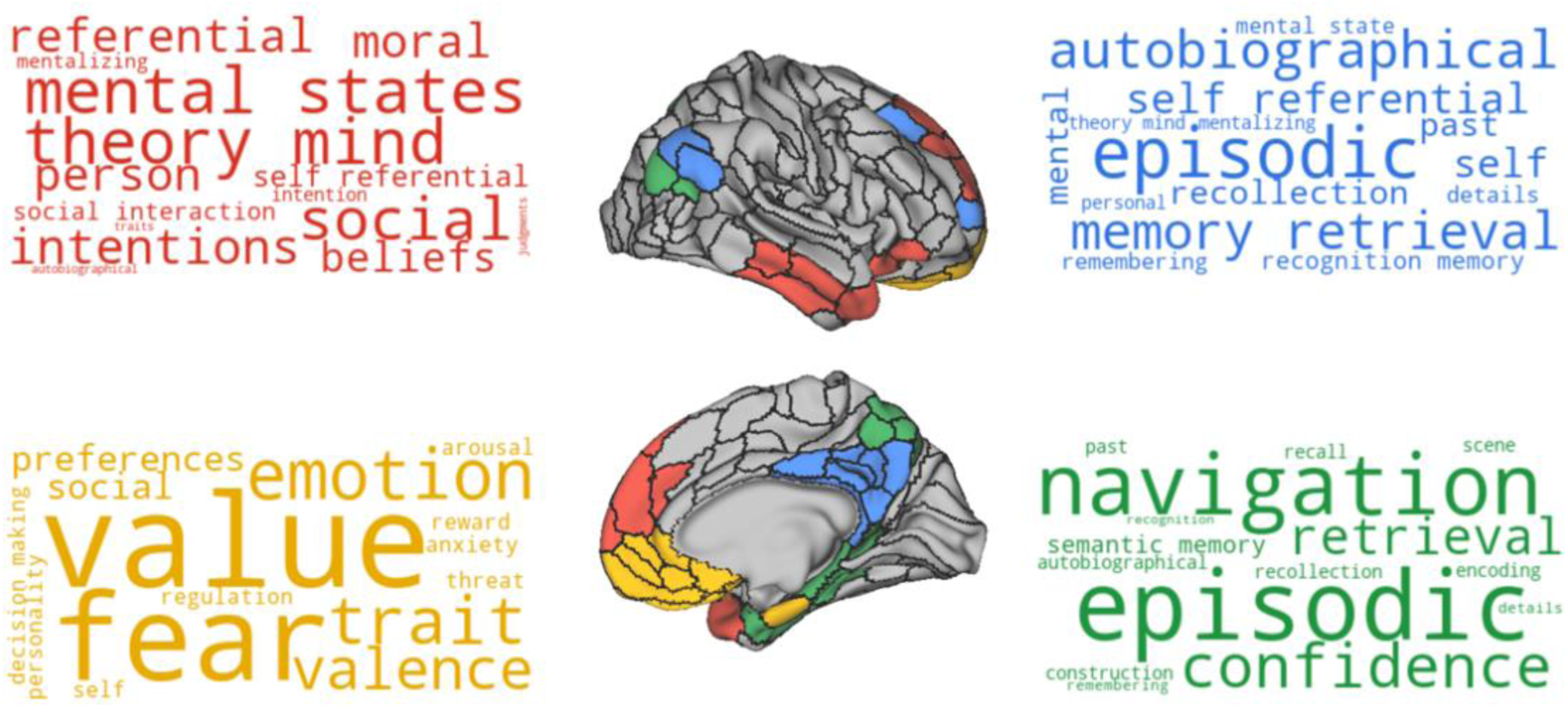
Terms associated with activation of each cortico-hippocampal network. Word clouds are colored according to the network they describe which are displayed on the inflated brain. The size of the terms in the word cloud relates to how strongly the meta-analytic activity of that term correlates with the spatial extent of the network. Red, AT subnetwork; Blue, PM subnetwork; Yellow, MP subnetwork; Green, Medial temporal network.

**Figure 8.**
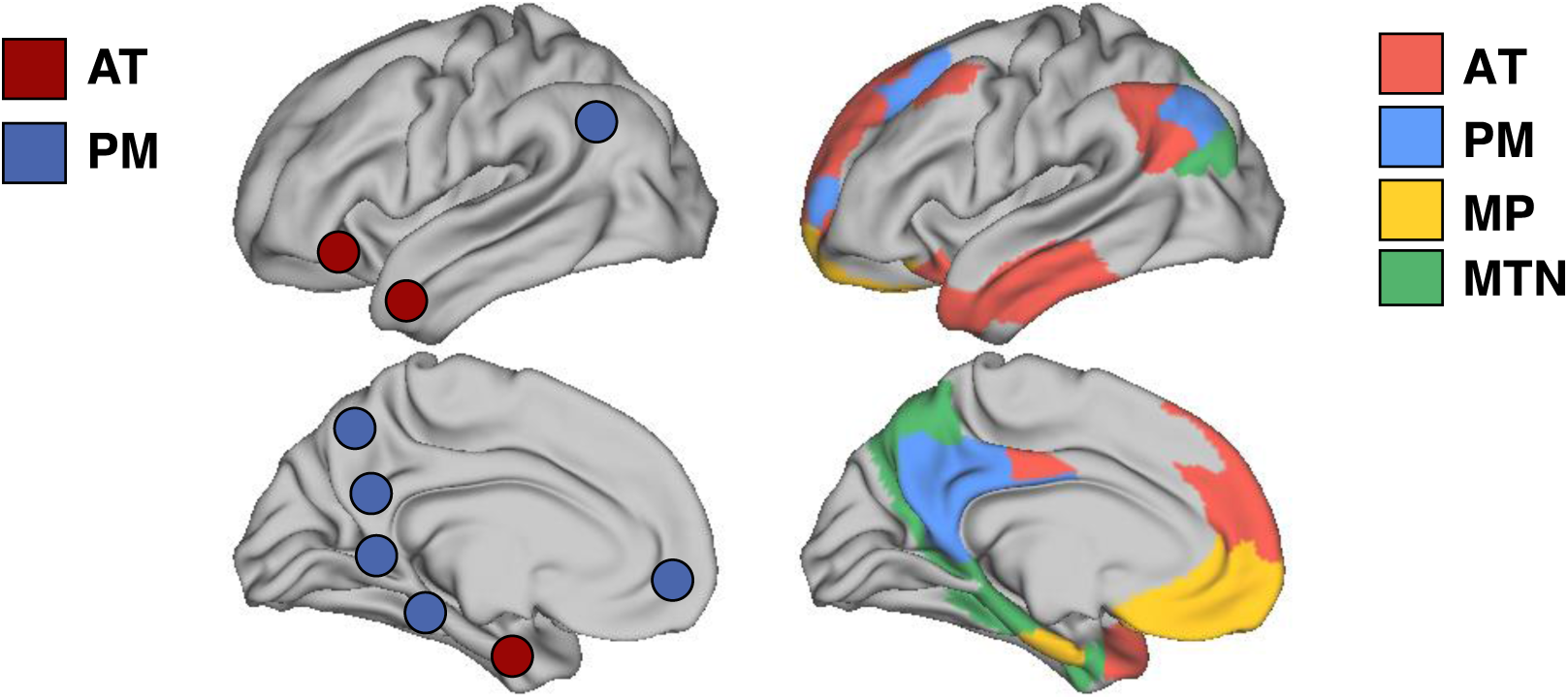
Left. Hypothesized PM/AT divisions with PM nodes as blue and AT nodes as red (29, 74). Right. Data-driven network assignments.

Although the present work shows that the MTN could be differentiated from the DMN, they do not imply that regions within the MTN are functionally or anatomically identical in terms of connectivity. Prior anatomical studies in animal models (75, 76) and fMRI studies in humans have shown that the PRC and PHC have different functional connectivity patterns (24–26, 77). In fMRI studies, these differences are most apparent when the functional connectivity maps of the PRC and PHC are directly contrasted to identify regions that show disproportionately high connectivity to the PRC or the PHC. When not contrasted, the two regions frequently show strong functional connectivity to each other and considerable overlap in the spatial extent of their functional connectivity (24, 25, 77). For example, Wang et al (25) and Zhuo et al. (77) showed that the PHC and posterior PRC show high connectivity with each other, and with medial parietal, medial prefrontal, lateral parietal and temporopolar regions. Therefore, the evidence presented here should not be taken to reflect the presence or absence of functional distinctions between PHC and PRC (as prior work has clearly shown that certain regions connect more to PHC or PRC), but rather they demonstrate that when taking into account the topology of functional connections across the entire neocortex, the MTN can be segregated from the DMN. This evidence suggests that heterogeneity exists within the previously reported PM network.

Treating the brain as an abstract graph, we observed that the MTN serves as a critical bridge between the visual network and the DMN and hippocampus, such that removal of the MTN results in considerable loss of cross-network connectivity between the visual network and DMN/hippocampus. The medial temporal cortex is thought to provide high-level feature representation of the external world to the hippocampus which then binds those representations in context (4, 22, 29, 58). Damage to the medial temporal cortex is often associated with varying level of circumscribed memory impairment. Our results here suggest that loss of MTN connections would result in disconnection between the external visual world and the hippocampus and DMN, which could result in memory impairment. Using a novel analytic method, a recent study mapped the functional connectivity of regions in the brain where lesion damage resulted in amnesia and found a common functional network to all the lesions (11). This lesion functional connectivity map encompassed the DMN, MTN and encroached into visual network regions as described in this study, underscoring the importance of these networks in memory processing. Another study of amnesia following traumatic brain injury observed reduced functional connectivity between the PHC and PCC was related to memory impairment and normalized when memory functioning improved (78).

The distinction between the MTN and the DMN is consistent with recent studies suggesting heterogeneity within the PM network (14, 38, 50, 79). Based on its network properties, the MTN sits in a reasonable position to represent higher-order features of the visual world to the DMN which is spatially distal from networks involved in primary sensation. Recent theories of event cognition have suggested a separation between content (higher order representations of current environment) and structure (abstracted knowledge pertaining the typical way in which sequences of events unfold) (80). One possible instantiation of this separation is that the MTN represents content of events whereas the DMN represents abstracted structure. Indeed, a recent study has shown that when orienting to retrieval of perceptual features during autobiographical recall, there is increased activity in the MTL, retrosplenial, and dorsal precuneus (regions in the MTN) compared to when recall is oriented to conceptual or thematic elements that preferentially resulted in activity of posterior cingulate, medial prefrontal, temporal polar and lateral parietal cortex (regions in the DMN) (39). Another study found a similar separation between the MTN and DMN by having participants encode a set of movie clips that were considered typical (events that one would be familiar with through day to day life or through movies/stories) or atypical (very unusual or dissimilar to anything you would encounter in day to day life or in movie/stories) (40). Regions in the DMN showed increased activity during encoding of typical compared to atypical events and the reverse was true for regions in the MTN. During encoding of typical events, activity in the DMN was interpreted as being related to retrieval of relevant prior event knowledge to assist in comprehension and integration of incoming information. Conversely, during encoding of atypical events, the activity in the MTN was interpreted to reflect the need to encode the relations between items, contexts and actions in the absence of a strong semantic framework. While these studies present strong evidence to suggest a distinction between the MTN and DMN, it is important to note that in many circumstances involving memory recall, these networks will be working in tandem to support retrieval of both content and structure with some degree of bias that may favor retrieval of item-level, scene-level or event-level elements. For example, one might expect that, even within the MTN, a contrast that focusses on the distinction between different types of content, such as scenes versus objects, may split regions within the MTN that have a bias towards those content types (35). These findings demonstrate the necessity of our intrinsic connectivity analyses to clarify that these differences in patterns of functional activity are likely due to the presence of distinct functional networks.

The second major finding in the present study is that the DMN could be differentiated into three subnetworks: the PM, AT, and MP subnetworks. The PM subnetwork consisted of posterior cingulate, angular gyrus, and dorsal prefrontal cortex, whereas the AT subnetwork consisted on temporopolar, lateral temporal, orbitofrontal and dorsal medial prefrontal cortex. The collection of regions in the PM and AT subnetwork are in line with the PM/AT framework (24, 25), with the notable exception that the parahippocampal, retrosplenial cortex were not in the PM subnetwork, and the perirhinal cortex was not in the AT subnetwork. The PM and AT subnetworks also resemble DMN subnetworks identified by previous data-driven methods (15, 41, 42), confirming that the PM and AT distinctions theorized do indeed persist, but they form distinct communities from their MTN counterparts when examining their connectivity architecture.

The MP subnetwork consists of medial prefrontal cortex, along with the entorhinal cortex and was not explicitly predicted by prior theories. The medial prefrontal cortex has long been distinguished from the orbitofrontal cortex based on anatomical connectivity (21, 22, 81), and is often grouped together with the posterior cingulate cortex as one of the “core hubs” of the DMN (15, 41, 42). The entorhinal cortex certainly has dense interconnections with the medial prefrontal cortex (81), but also is strongly interconnected with the hippocampus and adjacent medial temporal cortex (19, 22, 82). Interestingly, the entorhinal cortex often is part of the DMN even when the MTL cortex is distinguished from the rest of the DMN ((73, 83), but see (15, 47)). Furthermore, network analysis on histologically-defined axonal connections between cortical regions in rodents, showed that the lateral entorhinal cortex was part of a community with the prefrontal cortex, consistent with our partition, though the medial entorhinal cortex was placed in a community with the hippocampus, retrosplenial cortex and subiculum (84). A distinction in the functional connectivity of entorhinal cortex subregions has been shown in high-resolution human resting state data (23, 37), but the spatial resolution of the current study is unable to resolve these subregions and future work will be important to further characterize its connectivity in humans.

We examined the topology of how these subnetworks connect to each other and the rest of the brain finding that the MP subnetwork had the highest within-DMN participation coefficient, with nodes connecting broadly to AT and PM regions. Conversely, the MP subnetwork had a low out of DMN participation coefficient, especially relative to the AT subnetwork. Our results provide converging evidence for a recent study published by Gordon et al (85) that also found that a set of lateral networks resembling the AT subnetwork here, showed the highest participation coefficient out of network. Their AT equivalent showed intercommunication with language and frontoparietal networks, as was also seen here. The participation coefficient findings suggest that the AT subnetwork and several high participation PM nodes are positioned to mediate information transfer in and out of the DMN, whereas the MP subnetwork may integrate and coordinate information within the DMN. A recent review of the human lesion literature has reported that robust memory impairment can be observed following medial prefrontal cortex damage and suggested that this may be due to the fact that the medial prefrontal cortex has a role in initiating and coordinating cognitive processes for retrieval and episodic simulation (86). Given the network findings here, and implications from the lesion literature, the MP subnetwork is well positioned for higher-order integration and coordination of memory-guided activity.

Differentiation of the three DMN subnetworks was reinforced by evidence from an independent dataset in which we found that regions grouped in the same network represented similar kinds of information, as compared to regions in different networks. These findings replicate and extend results from previous studies that identified different functional differences between different cortico-hippocampal networks (17, 69). For instance, Ritchey et al. (17) used community detection to differentiate networks of regions hypothesized to show different patterns of MTL connectivity. Ritchey et al. (17) interrogated these networks in an independent study of episodic memory retrieval, as did Inhoff et al. (69) during a concept learning task. The functional distinction of these networks was supported by findings showing that regions grouped in the same network had greater representational profile similarity (17) and activation profile similarity than regions grouped in different networks during learning and memory tasks (17, 69). In these past studies, the regions used in network construction were those that showed a significant difference in functional connectivity between the PRC and PHC, being motivated by anatomical evidence suggesting parallel pathways in the MTL (19). Whereas the seed selection that defined these networks was based specifically on regions that had different functional connectivity between PHC and PRC, in the current study, we did not impose any pre-existing assumptions, and identified networks holistically in the context of the entire brain. We saw reliably higher representational profile similarity for regions in the same network—a pattern consistently found in every subject, highlighting the functional significance of these cortico-hippocampal networks and validating our delineation of these cortico-hippocampal networks.

We also found that the MTN and DMN subnetworks show variation in their connectivity across the anterior and posterior hippocampus. The hippocampus is known to vary along its long axis in terms of intrinsic connectivity, extrinsic connectivity, receptor distribution, and gene expression (19, 20, 87–90). In humans, resting state functional connectivity has been the primary tool for examining differences in connectivity along the hippocampal longitudinal axis, though distinctions between the anterior and posterior hippocampus have been observed using DTI (28). The anterior hippocampus has been shown to have greater connectivity to inferior temporal, temporopolar, orbitofrontal, medial prefrontal and PRC, whereas the posterior hippocampus has been shown to have greater connectivity to PHC, retrosplenial cortex, inferior parietal cortex, precuneus, the anterior thalamus (24, 26–28, 63). Here, we replicated and extended these findings to characterize hippocampal connectivity to the networks formed by these cortical regions. We found that the anterior hippocampus has relatively greater connectivity to the MP and AT subnetworks, whereas the posterior hippocampus has relatively greater connectivity to the MTN. The preferential connectivity to the anterior hippocampus was relatively consistent across all regions in the MP and AT subnetworks, but there was heterogeneity in the MTN’s preference for the anterior and posterior hippocampus with the anterior hippocampus having stronger connectivity to anterior MTL cortex and the posterior hippocampus having relatively stronger connectivity to dorsal medial parietal cortex. Theories regarding the functional difference of the anterior and posterior hippocampus are often informed by preferential long-axis connectivity to the rest of the brain (16, 91), but they do not consider how the rest of the brain connects to form networks. The results presented here suggest the possibility that functional differences between the anterior and posterior hippocampus may reflect differences in their connections to the subnetworks of the DMN.

To better understand the functional specializations of these networks, we used meta-analytic maps from the Neurosynth database to identify the cognitive constructs that have been reliably associated with regions in the DMN subnetworks. Activity within the MTN was associated with a set of terms heavily rooted in episodic memory and navigation. Activation in the MP subnetwork, in contrast, was highly associated with terms pertaining to emotion, and value. As noted above, the MTN (in particular the dorsal precuneus and dorsal posterior cingulate) had greater connectivity to the posterior relative to the anterior hippocampus, whereas the MP subnetwork had greater connectivity with the anterior compared to posterior hippocampus. These connectivity patterns and neural decoding results converge with theories linking the posterior hippocampus to cognitive and navigational processes and the anterior hippocampus in motivation and emotional behavior (92). We also found that activation in the AT subnetwork was associated with terms that relate to social cognition, theory of mind, and beliefs, in line with contemporary theories (93) and lesion evidence (94) of this network. In the PM subnetwork, Neurosynth decoding revealed cognitive terms related to episodic memory and the self. This is echoed by recent work, taking a highly subject-specific approach, that found a distinction within the DMN between regions involved in episodic versus theory of mind-like processes (95). In this study DiNicola et al (95) identified two distinct subnetworks of the DMN on a subject-by-subject, voxel-wise basis. One subnetwork resembled the AT subnetwork and the other resembled the PM subnetwork described here. The same subjects also performed both episodic projection tasks and theory of mind tasks and it was observed that the voxels classified as being in the AT-like network showed greater activity for the theory of mind tasks, whereas the voxels classified as being in the PM-like network showed greater activity for the episodic projection tasks. Interestingly, the Neurosynth analysis showed that the term “autobiographical” was strongly associated with all four cortico-hippocampal networks. This broad association may be due to the fact that autobiographical memory tasks usually involve retrieval of episodically rich events that dynamically unfold involving social interactions, and relevance to the self (96).

Although the Neurosynth decoding analysis is exploratory, and by definition, post-hoc, they can guide the generation of hypotheses to be tested in future studies. Further studies, similar to the work of DiNicola et al (95), should examine whether MTN and DMN subnetworks segregate based on spatial, valuation, and social processes, within the same group of individuals. For example, during encoding and recall of complex events, we might expect regions within the MTN to represent high-level object and context features, regions within the AT subnetwork to represent interpretations of theory of mind and social interactions, and regions within the MP subnetwork to represent the emotional valence or perceived value of the event. Indeed, a recent preprint demonstrated a dissociation between the MTN and MP subnetwork during event simulation, with increased activity in the MTN occurring when subjects were instructed to simulate past or future events using cues that were selected to elicit high levels of vividness, whereas increased MP activity was observed when cues elicited simulations with high levels of valence (97). Further, based on the functional connectivity of the MP subnetwork, we hypothesize that it would be well suited to serve a coordinating role in network rearrangement to accomplish task goals and demands. Again, a recent study by Nawa and Ando (98) found that when elaborating on autobiographical memories, the vmPFC drove hippocampal activity and this effect was augmented for memories that were of greater emotional valence. Further, damage to the medial prefrontal cortex is associated with memory impairments characterized by difficulties in elaborating on memories unfold over time and events (86). We hypothesize that these impairments may be associated with dysfunction of network dynamics due to a loss of the coordination normally provided by the MP subnetwork.

Finally, we note that recent studies have demonstrated the value in taking subject-specific approaches to delineating functional networks in a voxel-wise fashion (14, 47). This work has shown that there are individual differences in the location of boundaries between functional networks, though it is notable that the majority of the cortex is labelled consistently in the majority of participants (73). A key goal of the present study was to provide a clear framework to guide analyses in future task-fMRI studies, in which it might not be feasible to obtain independent, subject-specific parcellations. We have listed each region and its community label in Supplementary Table 3, and the HCP-MMP atlas is available online (https://balsa.wustl.edu/study/show/RVVG, (44)). Task fMRI studies that use group-level analyses can use the group-level characterization of cortico-hippocampal networks reported here in order to rigorously test hypotheses about the functions of these networks and how they may be implicated in memory disorders, as resting connectivity has shown to predict spread of pathology in Alzheimer’s (99), and DMN subnetwork perturbation has been demonstrated in MTL amnesia (100).

## Conclusions

The hippocampus affiliates with a broad set of regions that enable episodic retrieval. Here, we have shown that the MTN and three subnetworks of the DMN can be differentiated on the basis of their whole-brain functional connectivity. The three DMN subnetworks vary in connectivity along the hippocampal long axis, have distinct representational roles during a memory task, and topologically are connected by a set of hubs in the medial prefrontal subnetwork. This subnetwork organization offers a novel framework to investigate event cognition and memory retrieval.

## Methods

### Participants

Forty-five healthy, young adult participants were recruited from the University of California, Davis and surrounding area (N_Females_ = 26, mean age = 25.6 years [SD = 4.2 years]). All participants were right-handed and neurologically healthy. The study was approved by the Institutional Review Board of the University of California at Davis and all participants provided informed consent prior to participation. Participants were compensated $20/hour for their time. This sample size is comparable to the cohort sample sizes from the seminal Power et al. (101) study investigating functional brain organization. There is no specific effect size that can be taken to approximate power for identifying network communities, but previous studies have shown high reliability across subjects in network identification when using greater than 10 minutes of resting-state data (73, 102).

### MRI Acquisition

MRI scanning was performed using a 3T Siemens Skyra scanner system with a 32-channel head coil. A T1-weighted structural images was acquired using a magnetization prepared rapid acquisition gradient-echo pulse sequence (TR = 2100ms; TE = 2.98ms; field of view = 256 mm^2^; flip angle = 7°; image matrix = 256 × 256, 208 axial slices with 1.0 mm^3^ thickness; GRAPPA acceleration factor 2 with 24 reference lines). Functional images were acquired using a gradient EPI sequence (TR = 1220 ms; TE = 24 ms; field of view = 192 mm^2^ ; image matrix = 64 × 64; flip angle = 67°; bandwidth = 2442 Hx/pixel; multiband factor = 2; 38 interleaved axial slices, voxel size = 3 mm^3^ isotropic). Five runs of five minutes in duration were acquired for rest scans for a total of 25 minutes of resting state fMRI data per subject. Participants were instructed to lay as still as possible and try not to fall asleep.

### Anatomical Preprocessing

T1-weighted anatomical scans were preprocessed using FreeSurfer which included intensity normalization, removal of non-brain tissue, transformation to Talaraich space, and segmentation of gray matter, white matter, and CSF. Surfaces were calculated for the white matter-gray matter, and gray matter-pial interface. Surface-based registration to the HCP-MMP1.0 atlas (44) was performed and subject-specific cortical regions were calculated according to atlas boundaries. Surface-based cortical regions were converted to volumetric regions of interest and transformed into functional native space. The hippocampus was segmented in FreeSurfer in an automated fashion. Manual adjustments were done to correct misclassified voxels and the hippocampus was divided into anterior and posterior segments based off of the disappearance of the uncal notch (16), with the posterior hippocampus designated as all of the hippocampus posterior to the disappearance of the uncal notch on a coronal slice.

### Functional MRI Preprocessing

Functional preprocessing was performed using *fMRIPrep* 1.4.1 ((103, 104), RRID:SCR_016216), which is based on *Nipype* 1.2.0 ((105, 106), RRID:SCR_002502). For each of the 5 BOLD runs found per subject (across all tasks and sessions), the following preprocessing was performed. First, a reference volume and its skull-stripped version were generated using a custom methodology of *fMRIPrep*. A deformation field to correct for susceptibility distortions was estimated based on two echo-planar imaging (EPI) references with opposing phase-encoding directions, using 3dQwarp from Cox and Hyde (107) (AFNI 20160207). Based on the estimated susceptibility distortion, an unwarped BOLD reference was calculated for a more accurate co-registration with the anatomical reference. The BOLD reference was then co-registered to the T1w reference using flirt (FSL 5.0.9, (108)) with the boundary-based registration (109) cost-function. Co-registration was configured with nine degrees of freedom to account for distortions remaining in the BOLD reference. Head-motion parameters with respect to the BOLD reference (transformation matrices, and six corresponding rotation and translation parameters) are estimated before any spatiotemporal filtering using mcflirt (FSL 5.0.9, (110)). The BOLD time-series were resampled onto their original, native space by applying a single, composite transform to correct for head-motion and susceptibility distortions. These resampled BOLD time-series will be referred to as *preprocessed* BOLD. Several confounding time-series were calculated based on the *preprocessed BOLD* as part of *fMRIPrep*, however, only the head-motion estimates calculated in the correction step and frame displacement (FD) were used in subsequent analyses. All resamplings can be performed with *a single interpolation step* by composing all the pertinent transformations (i.e. head-motion transform matrices, susceptibility distortion correction when available, and co-registrations to anatomical space). Gridded (volumetric) resamplings were performed using antsApplyTransforms (ANTs), configured with Lanczos interpolation to minimize the smoothing effects of other kernels (111). Many internal operations of *fMRIPrep* use *Nilearn* 0.5.2 ((112), RRID:SCR_001362), mostly within the functional processing workflow. For more details of the pipeline, see https://fmriprep.readthedocs.io/en/stable/workflows.html.

The preprocessed BOLD timeseries, anatomical images, and native space Glasser parcels were imported into CONN Toolbox version 18b (www.nitrc.org/projects/conn, RRID:SCR_009550). Based on Ciric et al.’s (113) assessment of 14 common denoising protocols, we selected the ‘9p’ protocol because it was shown to facilitate the highest functional network identifiability (see (113) Figure 5b). BOLD timeseries were demeaned, linear and quadratic trends were removed, and bandpass filtered between .008 and .09 hz. The 9p protocol includes as confound regressors six motion parameters, WM, CSF, and global signal regression. All regressors were band-pass filtered to maintain the same frequency range as the data. Visual inspection of the frequency distributions of functional connectivity values as well as quality control plots of the correlation between mean FD and functional connectivity values was performed to identify any aberrant runs or subjects. Five outlier subjects were identified which each corresponded to previously identified subjects having a mean FD value of greater than .15mm. These subjects were excluded from further analyses. Functional connectivity matrices were created for each subject by computing Pearson’s correlations between all possible pairs of each regions’ confound-corrected timeseries. Finally, each correlation value was Fisher z-transformed with the inverse hyperbolic tangent function.

### Community Detection

Cortical functional connectivity (FC) matrices from all subjects were thresholded to exclude negative connections that may be introduced by global signal regression (114, 115) and then averaged together. Using the resulting group averaged connectivity matrix, community detection was performed using the Louvain method (45) via the brain connectivity toolbox (https://sites.google.com/site/bctnet/) which iteratively performs a greedy optimization of modularity by randomly selecting nodes and merging them into the community that maximally increases modularity, until no more gains in modularity are possible. Modularity (Q) describes how well a network can be divided into communities that have higher within-community connectivity than would be expected by chance and can range from −1 to 1, with negative values indicating fewer intracommunity connections than would be expected by chance and positive values indicating higher intracommunity connections than would be expected by chance (56). We used a method of the Louvain community detection algorithm adapted to accommodate weighted graphs (116) which accepts the weighted, group average functional connectivity matrix. This algorithm can be tuned using a resolution parameter, gamma, that biases the algorithm towards producing few, large networks (low gamma) or towards many, smaller networks (high gamma). To determine the resolution parameter in a principled manner, we adopted the approach used by Ji et al. (43). The criteria included: i) separation of the primary sensory-motor networks (visual, auditory, somatomotor) from all other networks, ii) high stability across nearby parameters (similar network partitions across neighboring parameter settings), and iii) optimized modularity.

Given that the Louvain algorithm detection is influenced by a random starting point, for every tested resolution parameter, we ran the algorithm 1000 times. To determine the optimal partitioning solution at each resolution, gamma, we examined how consistent a given partitioning solution was to every other solution produced over the 1000 iterations at the same resolution. Consistency was calculated using the z-Rand score (46) (http://commdetect.weebly.com/). For each partition solution, we weighted the mean z-Rand score (consistency) of the partition with the modularity value of that partition to select a partition solution that was both highly consistent at a given resolution and had high modularity. These methods are in keeping with Ji et al. (43). Using the default parameter, gamma = 1, we could not satisfy criteria i) as the somatomotor and auditory network were grouped together (also seen in Ji et al. (43)). Thus, we ran a parameter sweep from gamma 1 – 2.8 by increments of 0.005 until we identified a resolution parameter at which the somatomotor and auditory network separated consistently and produced the highest modularity-weighted z-Rand score.

### Path Length Analysis

Using the brain connectivity toolbox (https://sites.google.com/site/bctnet/), weighted path length was calculated between all nodes in the brain for each subject, after excluding negative connectivity weights. Path length is the fewest number of links that need to be traversed to connect two nodes. To calculate the weighted path length, the functional connectivity weights were inverted creating a distance matrix in which nodes with high functional connectivity had low distance. Then, the shortest path between each pair of nodes was calculated providing a value that represented the minimal the total weighted distance that needed to be traversed to connect two nodes.

We hypothesized that removal of the MTN would reduce the ability of the visual network and DMN/hippocampus to connect with each other. To test this hypothesis, all the nodes from the MTN were removed from each subject’s network and the average weighted path length was calculated between visual network and DMN and between visual network and hippocampus.

Because removal of nodes can only lead to an increase in path length (shorter paths can be removed, but will never be added following removal of nodes), we compared the weighted path length following removal of the MTN to the weighted path length following removal of every other network, using paired t-tests. As a secondary control, we also compared the weighted path length following removal of the MTN to the distribution of weighted path lengths following removal of 30 random nodes (other than the DMN, visual network and hippocampus) over 1000 iterations. For this, we calculated the z-score of the weighted path length following removal of the MTN compared to the distribution of weighted path lengths following removal of 30 nodes over 1000 iterations. This was done for each subject such that each subject had a corresponding z-score. A one-sample t-test was then performed on the z-score values to examine whether they were significantly greater than 0.

### Community Partitioning of the Default Mode Network

From the identified whole brain partitioning, we selected the default mode network based on visual similarity to previously identified DMN solutions (42, 43, 117). We repeated the community detection procedure described above on the identified DMN (1000 iterations of Louvain community detection at each resolution parameter) and calculated modularity-weighted z-rand scores across a range of resolution parameters, 0.75 – 1.1 at increments of 0.05. We selected local modularity-weighted z-rand peaks as solutions for further exploration.

### Connection Diversity in the Default Mode Network

The participation coefficient, which quantifies the degree to which a node is connected to a diverse set of communities, was calculated using the tools from the brain connectivity toolbox (https://sites.google.com/site/bctnet/). Since the participation coefficient calculation uses node strength (the sum of the connectivity weights across the network) in the denominator of its calculation, nodes with unusually low strength can produce unstably large participation coefficient values. We, thus, excluded regions with node strength in the bottom 25% of the network hubs. Participation coefficient calculations were performed across a range of network densities from 5-20% at 1% intervals. These methods for calculating participation coefficient are in line with best practices in the literature (57, 59, 62, 118, 119). Whole network participation coefficient was calculated for each DMN region, using the whole brain connectivity matrix and community labels generated from the whole-brain Louvain community detection. Within DMN participation coefficient was also calculated for each DMN node, using only the connections within the DMN and using the DMN subnetwork labels. Participation coefficient values were compared between DMN subnetworks using a within-subjects mixed model using mixedlm from statsmodels in Python 3 (120).

### Hippocampal-cortical network connectivity

To identify hippocampal-cortical networks we calculated functional connectivity between each hippocampal ROI and every cortical ROI in the brain. To identify average connectivity of the hippocampus to a given network within-subject, functional connectivity weights were averaged together across cortical nodes, based on community affiliation. A one-sample t-test was performed to determine what networks were, on average, significantly connected to the hippocampus. Significance was set at *p*<.05, Bonferroni corrected. We examined the connectivity of the resulting significant networks and subnetworks to the hippocampus as a function of the long-axis (anterior versus posterior) and hemisphere (left versus right) in a within-subjects model as described above.

ROI level functional connectivity difference between the anterior and posterior hippocampus were performed in CONN toolbox version 18b. Using the cortical ROIs that were a part of large-scale networks connected to the hippocampus, a within-subjects analysis was performed which contrasted functional connectivity in the left and right anterior hippocampus against the left and right posterior hippocampus. Significance was set at *p*<.05, *FWE* corrected.

### Representational Similarity Analysis

Using an independent dataset collected during a memory-based decision-making task (68), we examined the representational profile of each ROI in the DMN subnetworks (See SI Appendix for a detailed description of the task). If the hippocampal-cortical networks identified using resting-state functional connectivity are functionally meaningful, then regions within those networks should have similar representational profiles.

In brief, 22 participants in this dataset viewed a set of 8 grocery items one at a time, in a pre-scanning session, and learned through trial and error whether each grocery item was desirable to a hypothetical customer within a store context or not. In each trial, a grocery item appeared in the context of a grocery store. There were four different grocery stores and for half of the items the particular grocery store modulated their desirability and for the other half, the grocery store context was irrelevant to their desirability. For example, the apple at grocery store ‘A’ may be desirable, but the apple at grocery store ‘B’ may be undesirable which makes apple’s desirability context-dependent. Alternatively, desirability of a carton of milk may be the same regardless or grocery store which makes it context-invariant. Overall preference averaged to 50% desirable for both context dependent and invariant. Further, of the four contexts, 50% shared the same preference rules. Following successful learning, participants were put in the scanner and shown the context and a food item, one at a time and asked to remember whether the food item was desirable to the hypothetical customer at the store context based on their learning phase.

Feedback was not provided during scanning. In the original study, Mizrak et al. (68) examined representational similarity between memory-based decisions depending on a) shared features between grocery items such as being modulated by the context or not, b) shared features between store contexts such as having similar desirability for the same grocery items or not.

Here, using the RSA toolbox (https://www.mrc-cbu.cam.ac.uk/methods-and-resources/toolboxes/license/, (121)), pattern similarity was calculated between memory based decision making trials for each ROI using Pearson’s correlation, excluding a) trials that occurred during the same scanning run, and b) incorrect trials. Thus, fluctuations in pattern similarity would be driven by variations in shared features. Correlations were then calculated between ROIs to calculate the similarity of their representational profiles. We then examined the representational similarity within cortico-hippocampal subnetworks versus between using a paired t-test to examine whether these networks identified using resting-state functional connectivity did indeed create functionally relevant communities.

To account for univariate effects on the multivariate analysis, we further calculated the mean BOLD activity within an ROI for each trial and correlated the trial-wise mean BOLD activity for each pair of ROIs to create an ROI-by-ROI activation similarity matrix. Using the activation similarity matrix, we regressed out the influence of univariate activity from the RDM similarity matrix and re-ran the statistical analysis contrasting within- vs. between-network RDM similarity. To account for the effects of spatial proximity between ROIs, we calculated the Euclidean distance between the centroids of each pair of ROIs to create an ROI-by-ROI distance matrix. Using the distance matrix, we regressed out the influence of spatial proximity from the RDM similarity matrix and re-ran the statistical analysis contrasting within- vs. between-network RDM similarity.

### Cognitive characterization of hippocampal-cortical networks

We decoded cortico-hippocampal networks to determine what terms are most frequently descriptive of the spatial distribution of the networks using the repository data version 0.7 (https://github.com/neurosynth/neurosynth-data) of Neurosynth (https://github.com/neurosynth/neurosynth) (70). The meta-analytic activation of every term in the Neurosynth database was correlated with the binarized volumetric masks of the hippocampal-cortical networks. For each hippocampal-cortical network we examined the top terms whose meta-analytic activation correlated most strongly with the spatial layout of the hippocampal-cortical network. We removed terms that related to anatomy (e.g. ventromedial prefrontal cortex), network (e.g. default mode), technique (e.g. independent component) and report terms associated with cognitive functioning (e.g. navigation). We also removed terms that were redundant with other terms (e.g. “autobiographic” and “autobiographical memory”). When removing redundant terms, the term that was more specific was retained (e.g. “autobiographical memory” retained in favor of “memory”), otherwise the term with the higher correlation to the network was retained. Word cloud visualizations were made using wordcloud 1.7.0 (https://pypi.org/project/wordcloud/).

