## Supplemental Information for "Organization of cortico-hippocampal networks in the human brain"

Supplemental Table 1. Mean of the average functional connectivity between the hippocampus and the regions within each cortical network across subjects with standard deviation in parenthesis. Functional connectivity represented as the Fisher z-transformed correlation between the functional timeseries between two regions.

| Network | Right Posterior | Left Posterior | Right Anterior | Left Anterior |
|---|---|---|---|---|
| Visual | -0.005 (.07) | -.027 (.07) | -.013 (.07) | -.031 (.08) |
| Somatomotor | .038 (.07) | .022 (.07) | **.063 (.08)** | **.052 (.09)** |
| Cingulo-Opercular 1 | -.021 (.06) | -.020 (.06) | -.089 (.07) | -.069 (.08) |
| Cingulo-Opercular 2 | -.091 (.1) | -.098 (.11) | -.084 (.12) | -.081 (.1) |
| DAN 1 | -.012 (.11) | -.042 (.11) | -.097 (.14) | -.084 (.14) |
| DAN 2 | -.053 (.06) | -.066 (.06) | -.073 (.07) | -.081 (.07) |
| Language | -.016 (.04) | -.004 (.05) | **.041 (.05)** | **.041 (.05)** |
| Frontoparietal | -.043 (.04) | -.032 (.04) | -.034 (.05) | -.034 (.04) |
| Auditory | .03 (.07) | .025 (.08) | .020 (.08) | .032 (.08) |
| DMN | **.082 (.05)** | **.092 (.05)** | **.111 (.06)** | **.118 (.06)** |
| MTL | **.056 (.04)** | **.038 (.04)** | **.042 (.05)** | **.028 (.04)** |

DAN, Dorsal Attention Network; DMN default mode network; MTL, medial temporal lobe network; Bold values are significant at $p<.05$, Bonferroni-corrected.

Supplemental Table 2. Correlation values of spatial overlap between the cortico-hippocampal networks and meta-analytic maps for cognitive terms.

| Cognitive Term | MTN | PM | AT | MP |
|---|---|---|---|---|
| episodic | 0.2009 | 0.1455 | 0.0634 | 0.0366 |
| navigation | 0.1499 | -0.0038 | -0.0462 | -0.0413 |
| confidence | 0.1479 | 0.0668 | 0.0169 | 0.0161 |
| retrieval | 0.144 | 0.0282 | 0.0576 | 0.0078 |
| semantic memory | 0.1425 | 0.0719 | 0.0615 | 0.0422 |
| autobiographical | 0.1411 | 0.1427 | 0.1231 | 0.096 |
| recollection | 0.1375 | 0.1062 | 0.0727 | 0.0212 |
| construction | 0.1289 | 0.067 | 0.0763 | 0.0398 |
| encoding | 0.1268 | 0.0058 | -0.011 | -0.0354 |
| scene | 0.119 | 0.0076 | 0.0323 | -0.0046 |
| past | 0.1175 | 0.1026 | 0.0863 | 0.068 |
| recall | 0.1133 | 0.0732 | 0.0574 | 0.0201 |
| remembering | 0.1029 | 0.0915 | 0.0571 | 0.0379 |
| details | 0.1013 | 0.0892 | 0.0356 | 0.0409 |
| recognition | 0.0991 | 0.0526 | 0.0177 | -0.0298 |
| mental states | 0.0377 | 0.0659 | 0.2203 | 0.047 |
| theory mind | 0.0401 | 0.0785 | 0.2154 | 0.0631 |
| social | 0.0394 | 0.051 | 0.188 | 0.0379 |
| intentions | 0.0263 | 0.0193 | 0.1696 | 0.0232 |
| referential | 0.0616 | 0.1285 | 0.1479 | 0.0995 |

| | | | | |
|---|---|---|---|---|
| person | 0.0529 | 0.0511 | 0.1473 | 0.0688 |
| moral | 0.0246 | 0.0729 | 0.147 | 0.0906 |
| beliefs | 0.0246 | 0.0514 | 0.1454 | 0.0521 |
| self referential | 0.061 | 0.1315 | 0.1454 | 0.0961 |
| social interaction | 0.0028 | -0.0078 | 0.1382 | 0.0379 |
| mentalizing | 0.0555 | 0.0788 | 0.1381 | 0.0544 |
| intention | 0.0382 | 0.0662 | 0.1242 | 0.0529 |
| traits | -0.0141 | 0.0039 | 0.1224 | 0.1244 |
| judgments | 0.0527 | 0.071 | 0.1109 | 0.0274 |
| memory retrieval | 0.1223 | 0.1323 | 0.0887 | 0.0514 |
| self | 0.061 | 0.1131 | 0.1404 | 0.1266 |
| mental | 0.0732 | 0.1061 | 0.1219 | -0.0033 |
| recognition memory | 0.0735 | 0.1009 | 0.0253 | -0.0232 |
| mental state | 0.081 | 0.083 | 0.1306 | 0.04 |
| personal | 0.0937 | 0.0755 | 0.1057 | 0.1002 |
| value | 0.0068 | 0.0321 | 0.0515 | 0.1706 |
| fear | -0.0059 | -0.0284 | 0.0041 | 0.1591 |
| emotion | 0.0027 | -0.0032 | 0.0987 | 0.156 |
| trait | -0.0109 | 0.0306 | 0.1134 | 0.1422 |
| valence | 0.0124 | 0.0052 | 0.0861 | 0.1415 |
| preferences | -0.0208 | 0.0059 | 0.0209 | 0.13 |
| social | 0.0394 | 0.0378 | 0.188 | 0.1298 |

| | | | | |
|---|---|---|---|---|
| regulation | -0.0061 | 0.0044 | 0.0763 | 0.1284 |
| decision making | -0.0433 | 0.0404 | 0.0035 | 0.1272 |
| reward | -0.0322 | -0.0081 | -0.0101 | 0.1272 |
| personality | -0.0062 | 0.0254 | 0.1044 | 0.1267 |
| threat | 0.0107 | -0.023 | -0.0017 | 0.1234 |
| anxiety | 0.0179 | 0.0065 | 0.0271 | 0.1231 |
| arousal | -0.0055 | -0.0114 | 0.0389 | 0.1172 |

Supplemental Table 3. Regions of interest in the HCP-MMP atlas 1.0 that are found in the Medial Temporal network (MTN), Anterior temporal (AT) subnetwork, Posterior medial (PM) subnetwork, and Medial Prefrontal (MP) Subnetwork, with their community label provided.

| Region of Interest | Community | Name |
|---|---|---|
| L_POS2_ROI | MTN | Parieto-Occipital_Sulcus_Area_2 |
| L_PCV_ROI | MTN | PreCuneus_Visual_Area |
| L_7Pm_ROI | MTN | Medial Area 7P |
| L_7Am_ROI | MTN | Medial Area 7A |
| L_7Pl_ROI | MTN | Lateral Area 7P |
| L_PreS_ROI | MTN | Presubiculum |
| L_ProS_ROI | MTN | ProStriate Area |
| L_PeEc_ROI | MTN | Perirhinal Ectorhinal Cortex |
| L_PHA1_ROI | MTN | Parahippocampal Area 1 |
| L_PHA3_ROI | MTN | Parahippocampal Area 3 |
| L_TPOJ3_ROI | MTN | Area TemporoParietoOccipital Junction 3 |
| L_DVT_ROI | MTN | Dorsal Transitional Visual Area |
| L_PGp_ROI | MTN | Area PGp |
| L_PHA2_ROI | MTN | Parahippocampal Area 2 |
| R_POS2_ROI | MTN | Parieto-Occipital Sulcus Area 2 |
| R_PCV_ROI | MTN | PreCuneus Visual Area |
| R_7Pm_ROI | MTN | Medial Area 7P |
| R_POS1_ROI | MTN | Parieto-Occipital Sulcus Area 1 |
| R_7Am_ROI | MTN | Medial Area 7A |

| | | |
|---|---|---|
| R_7PL_ROI | MTN | Lateral Area 7P |
| R_PreS_ROI | MTN | Presubiculum |
| R_ProS_ROI | MTN | ProStriate Area |
| R_PeEc_ROI | MTN | Perirhinal Ectorhinal Cortex |
| R_PHA1_ROI | MTN | Parahippocampal Area 1 |
| R_PHA3_ROI | MTN | Parahippocampal Area 3 |
| R_TF_ROI | MTN | Area TF |
| R_TPOJ3_ROI | MTN | Area TemporoParietoOccipital Junction 3 |
| R_PGp_ROI | MTN | Area PGp |
| R_IP0_ROI | MTN | Area IntraParietal 0 |
| R_PHA2_ROI | MTN | Parahippocampal Area 2 |
| L_23d_ROI | AT | Area 23d |
| L_d32_ROI | AT | Area dorsal 32 |
| L_8Av_ROI | AT | Area 8Av |
| L_9m_ROI | AT | Area 9 Middle |
| L_8BL_ROI | AT | Area 8B Lateral |
| L_9p_ROI | AT | Area 9 Posterior |
| L_10d_ROI | AT | Area 10d |
| L_9a_ROI | AT | Area 9 anterior |
| L_a10p_ROI | AT | Area anterior 10p |
| L_47s_ROI | AT | Area 47s |
| L_TGd_ROI | AT | Area TG dorsal |
| L_TE1a_ROI | AT | Area TE1 anterior |
| L_TE2a_ROI | AT | Area TE2 anterior |
| L_PFm_ROI | AT | Area PFm Complex |
| L_PGi_ROI | AT | Area PGi |
| L_STSva_ROI | AT | Area STSv anterior |

| | | |
|---|---|---|
| L_TE1m_ROI | AT | Area TE1 Middle |
| R_d32_ROI | AT | Area dorsal 32 |
| R_9m_ROI | AT | Area 9 Middle |
| R_8BL_ROI | AT | Area 8B Lateral |
| R_9p_ROI | AT | Area 9 Posterior |
| R_10d_ROI | AT | Area 10d |
| R_47l_ROI | AT | Area 47 lateral |
| R_9a_ROI | AT | Area 9 anterior |
| R_47s_ROI | AT | Area 47s |
| R_TGd_ROI | AT | Area TG dorsal |
| R_TE1a_ROI | AT | Area TE1 anterior |
| R_TE2a_ROI | AT | Area TE2 anterior |
| R_STSva_ROI | AT | Area STSv anterior |
| R_TE1m_ROI | AT | Area TE1 Middle |
| L_RSC_ROI | PM | RetroSplenial Complex |
| L_7m_ROI | PM | Area 7m |
| L_POS1_ROI | PM | Parieto-Occipital Sulcus Area 1 |
| L_v23ab_ROI | PM | Area ventral 23 a+b |
| L_d23ab_ROI | PM | Area dorsal 23 a+b |
| L_31pv_ROI | PM | Area 31p ventral |
| L_8Ad_ROI | PM | Area 8Ad |
| L_s6-8_ROI | PM | Superior 6-8 Transitional Area |
| L_PGs_ROI | PM | Area PGs |
| L_31pd_ROI | PM | Area 31pd |
| L_31a_ROI | PM | Area 31a |
| L_p10p_ROI | PM | Area posterior 10p |
| R_RSC_ROI | PM | RetroSplenial Complex |

| | | |
|---|---|---|
| R_7m_ROI | PM | Area 7m |
| R_23d_ROI | PM | Area 23d |
| R_v23ab_ROI | PM | Area ventral 23 a+b |
| R_d23ab_ROI | PM | Area dorsal 23 a+b |
| R_31pv_ROI | PM | Area 31p ventral |
| R_8Ad_ROI | PM | Area 8Ad |
| R_PGi_ROI | PM | Area PGi |
| R_PGs_ROI | PM | Area PGs |
| R_31pd_ROI | PM | Area 31pd |
| R_31a_ROI | PM | Area 31a |
| R_p10p_ROI | PM | Area posterior 10p |
| L_a24_ROI | MP | Area a24 |
| L_p32_ROI | MP | Area p32 |
| L_10r_ROI | MP | Area 10r |
| L_47m_ROI | MP | Area 47m |
| L_10v_ROI | MP | Area 10v |
| L_10pp_ROI | MP | Polar 10p |
| L_OFC_ROI | MP | Orbital Frontal Complex |
| L_EC_ROI | MP | Entorhinal Cortex |
| L_25_ROI | MP | Area 25 |
| L_s32_ROI | MP | Area s32 |
| L_pOFC_ROI | MP | posterior OFC Complex |
| R_a24_ROI | MP | Area a24 |
| R_p32_ROI | MP | Area p32 |
| R_10r_ROI | MP | Area 10r |
| R_47m_ROI | MP | Area 47m |
| R_10v_ROI | MP | Area 10v |
| R_10pp_ROI | MP | Polar 10p |
| R_OFC_ROI | MP | Orbital Frontal Complex |

| | | |
|---|---|---|
| R_EC_ROI | MP | Entorhinal Cortex |
| R_25_ROI | MP | Area 25 |
| R_s32_ROI | MP | Area s32 |
| R_pOFC_ROI | MP | posterior OFC Complex |

S1 Figure Caption
Boxplots displaying the path length between the DMN and each of the other networks (targets) following removal of a given non-target network.

S2 Figure Caption
Boxplots displaying the path length between the hippocampus and each of the other networks (targets) following removal of a given non-target network.

S1 Figure

X-axis = removed network
Y-axis = path length between DMN and target
following removal of network

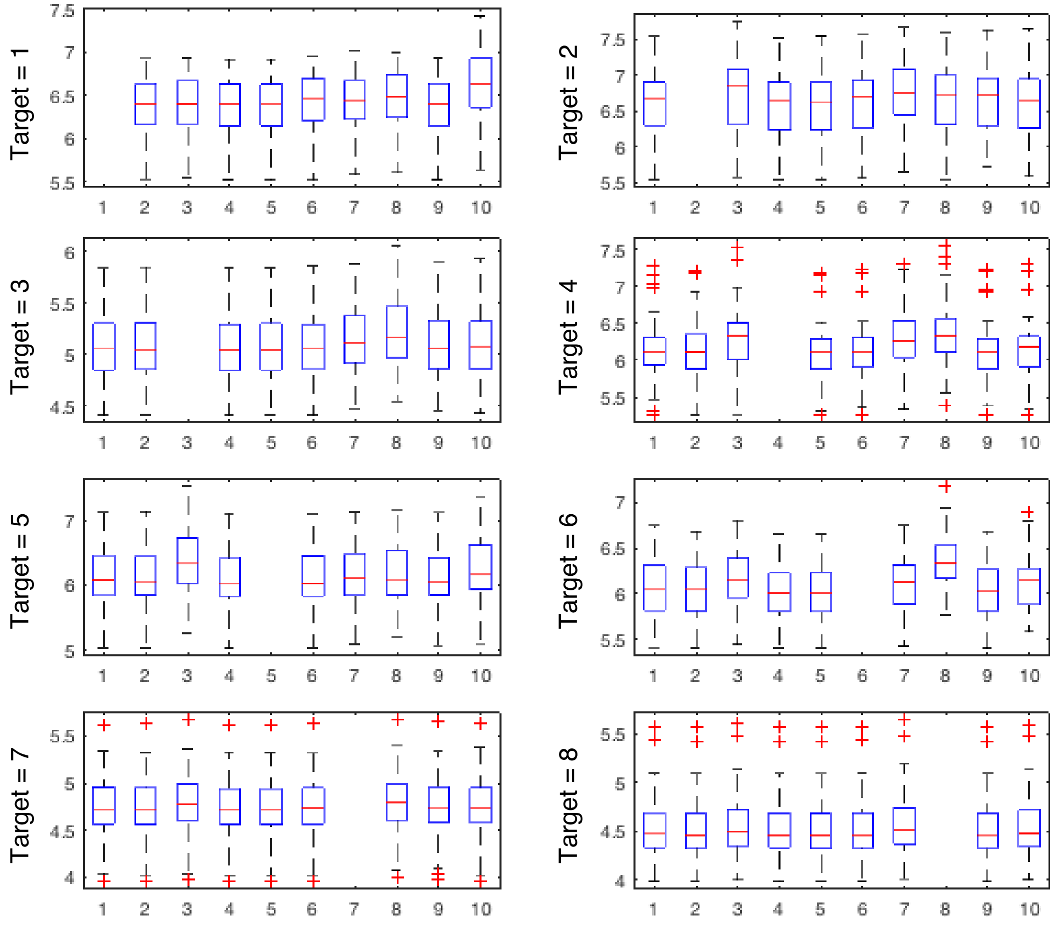

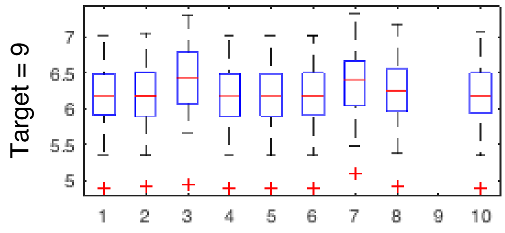

Labels

| | | | |
|---|---|---|---|
| 1 | Visual | 6 | Dorsal Attn 2 |
| 2 | somatomotor | 7 | Language |
| 3 | Salience 1 | 8 | Frontoparietal |
| 4 | Salience 2 | 9 | Auditory |
| 5 | Dorsal Attn 1 | 10 | MTN |

S2 Figure

X-axis = removed network
Y-axis = path length between hippocampus and target following removal of network

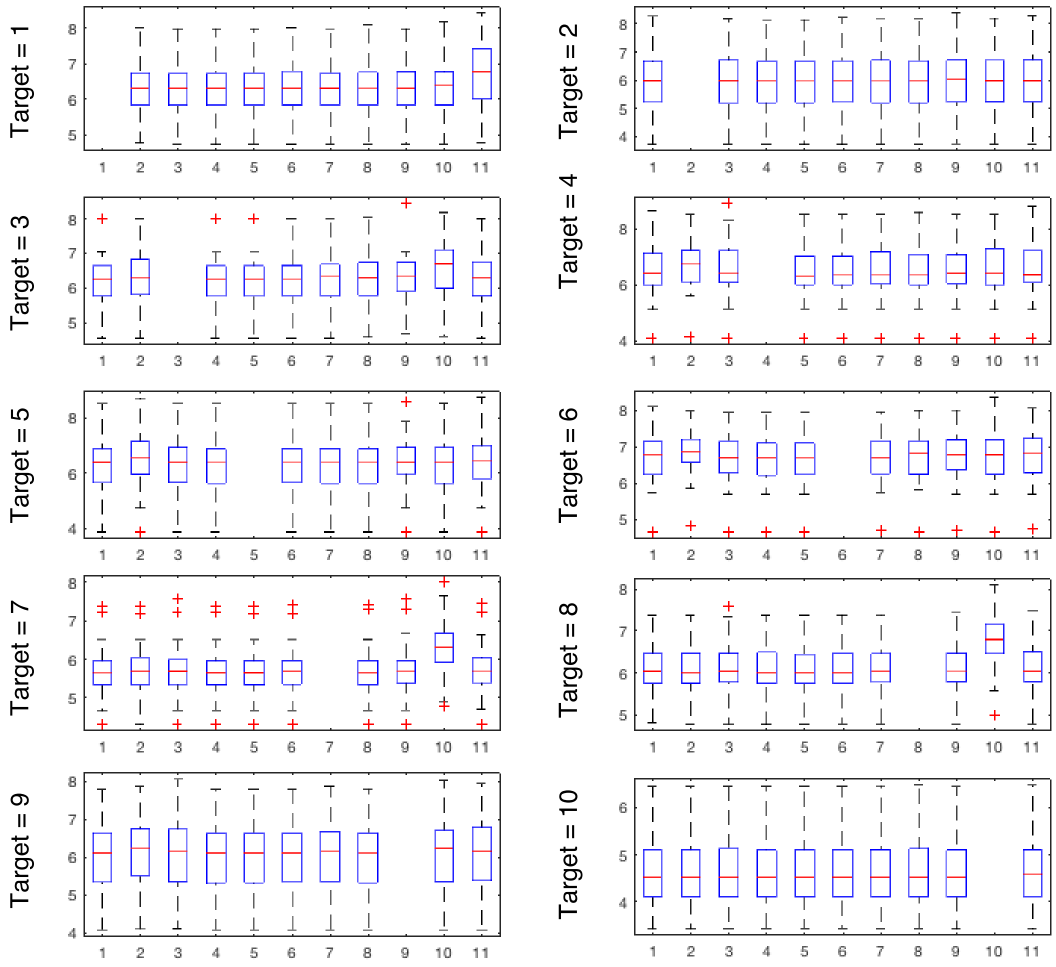

Labels

| | | | | | |
|---|---|---|---|---|---|
| 1 | Visual | 6 | Dorsal Attn 2 | 11 | MTN |
| 2 | somatomotor | 7 | Language | | |
| 3 | Salience 1 | 8 | Frontoparietal | | |
| 4 | Salience 2 | 9 | Auditory | | |
| 5 | Dorsal Attn 1 | 10 | DMN | | |
